# A urine-dependent human urothelial organoid offers a promising alternative to rodent models of infection

**DOI:** 10.1101/152033

**Authors:** Harry Horsley, Dhanuson Dharmasena, James Malone-Lee, Jennifer L. Rohn

## Abstract

Murine models describe a defined host/pathogen interaction for urinary tract infection, but human cell studies are scant. Although recent human urothelial organoid models are promising, none demonstrate long-term tolerance to urine, the natural substrate of the tissue and of the uropathogens that live there. We developed a novel human organoid from progenitor cells which demonstrates key structural hallmarks and biomarkers of the urothelium. After three weeks of transwell culture with 100% urine at the apical interface, the organoid stratified into multiple layers. The apical surface differentiated into enlarged and flattened umbrella-like cells bearing characteristic tight junctions, structures resembling asymmetric unit membrane plaques, and a glycosaminoglycan layer. The apical cells also expressed apical cytokeratin-20, a spatial feature of the mammalian urothelium. Urine itself was necessary for full development, and undifferentiated cells were urine-tolerant despite the lack of membrane plaques and a glycosaminoglycan layer. Infection with *Enterococcus faecalis* revealed the expected invasive outcome, including urothelial sloughing and the formation of intracellular colonies similar to those previously observed in patient cells. This new biomimetic model could help illuminate invasive behaviours of uropathogens, and serve as a reproducible test bed for disease formation, treatment and resolution in patients.

## Introduction

UTIs are amongst the most common infectious diseases worldwide, but despite being associated with substantial economic and human cost^1,2^, they are grossly understudied relative to other human diseases. UTI pathogens are also of particular concern in the global antibiotic resistance crisis, so their burden will only increase in the future^3^. Recurrence of infection even after antibiotic treatment is a particularly troublesome aspect of UTI, usually involving the same strain implicated in the first infection^1,4^. For example, among healthy young women who suffer from their first UTI, the risk of recurrence within 6 months is 24%^2^. If they had a history of one or more UTIs, the likelihood of recurrence rises to 70% in that same year ^2^. In another study, 14% of the 30,851 residents with UTI had more than one episode during the two-year study period, and 2% had six or more episodes^5^. These findings suggest that current treatment regimens are not ideal.

UTI is also problematic in more vulnerable subgroups: the risk of UTI dramatically increases in people with multiple sclerosis (MS)^6,7^, spinal injury^8^, renal transplant patients^9^ and anyone requiring urinary catheterization or other indwelling devices^10^. In fact, UTIs accounted for 10.5 million ambulatory care visits in 2007 in the United States^11^, with the direct and indirect costs being estimated at more than US$3.5 billion. This is likely to rise with the ageing global population and the emerging threat of antibiotic resistance. Finally, amongst the elderly, UTIs are one of the most commonly diagnosed infections^12^. More frequent UTI in these cohorts is not merely bothersome; UTI is known to exacerbate MS^13^, lead to confusion and falls in the elderly^14^, and increase the risk of organ rejection in renal transplant patients^15^. Furthermore, catheter-associated UTI carries an increased risk of urosepsis^16^, and bacteriuria in pregnant women is associated with preterm birth and other maternal morbidities^17^.

To understand why urinary infections are often recalcitrant to treatment, the pathogens must be studied in their unique environment. The urinary bladder is lined by a specialised transitional urothelium comprising 3-7 layers of cells: basal cells (above the basement membrane), intermediate cells (above basal cells) and morphologically distinct, highly specialised, often binucleated umbrella cells at the apical surface, which face outward into the bladder lumen^18^. These enlarged, flattened urothelial umbrella cells (or ‘facet cells’) partition urine and are thought to act as a powerful barrier to protect underlying tissue from harmful waste compounds^19^. They elaborate a highly-durable apical asymmetric unit membrane (AUM) consisting of thousands of regularly arrayed plaques or ‘facets’ approximately 16.5 nm across made up of four mannosylated transmembrane glycoproteins called uroplakins (UP)^19-21^.

In addition to the uroplakin family, the urothelium also elaborates a mucopolysaccharide-rich layer of glycosaminoglycans (GAG) which is believed to protect the bladder from infection and urine-born irritants^22^, of which chondroitin sulphate, heparan sulphate, hyaluronic acid, dermatan sulphate and keratin sulphate are the most studie ^23^. Chondroitin sulphate, in particular, is believed to play a key role in urothelial barrier function and exhibits luminal and basal expression in both human and porcine bladders^24^. In contrast, only heparan sulphate was detected in the luminal portion of calf bladders, elucidating possible differences between species^25^.

A significant proportion of research on the urothelium has been conducted using mouse models^21,26^. These findings have been widely translated into human oncology to locate the primary origin of metastatic tumours^27^ and to understand the biology of UTI^26^. While invaluable in many cases and necessary for regulatory approval of drugs, some animal models of human disease, the majority of which are murine, have received widespread criticism in recent years^28-32^. The limitations of murine models are particularly evident when modelling human infection and attempting to treat this induced pathology with novel antimicrobials^28^. In such studies, mice are frequently infected with far higher quantities of log-phase bacteria than would be evident in a slow-growing chronic human infection, and the pharmacokinetic profiles of a given drug are challenging to translate to humans^28,33^.

In the case of urinary infection studies, it is known that the human and mouse bladder urothelium differ in a number of structural ways. The markers expressed are similar, but in contrast to the murine model, human bladder urothelial marker expression exhibits a relationship with the level of cellular differentiation^21,34^. For example, the healthy human bladder has been shown to express cytokeratin 20 at the luminal surface whereas cytokeratin 8 is expressed throughout the cells of the urothelium^18,35^. The incorrect spatial expression of cytokeratin 20 by terminally differentiated umbrella cells has been linked to painful bladder syndrome and neurogenic bladder and is thought to predispose people with MS to chronic UTI^6,18,35^. Studies also suggest that murine and human bladders can differ in their innate immune response to uropathogens (for example in their expression and use of Toll-like receptors^36^). Moreover, rodent bladders differ from those of humans functionally. While larger mammals (>3Kg) share a scalable urinary capacity and consistent voiding duration, rodents urinate almost constantly, bringing into question whether their bladders are a true storage organ^37^. The multiple disparities between the rodent and human bladders raise the possibility that relying so heavily on the former could be problematic for understanding UTI in the latter.

Given these species differences, there is a need for alternative human-based models to augment the impressive body of elegant *in vivo* mouse experiments into UTI biology. Human bladder cancer cell lines grow readily and are tractable, but they are genetically abnormal and, therefore, bear little resemblance to primary urothelial cells in terms of structure and function. In particular, although some retain the ability to form a stratified organoid, they do not form an organised and differentiated 3D architecture^38,39^, which is crucial not least for understanding host/pathogen interaction, as uropathogens are proposed to invade the urothelium via binding to factors only present in terminally differentiated umbrella cells^26^.

On the other hand, recent years have seen advances in three-dimensional tissue engineering. A number of promising 3D urothelial models have been described in the literature, the majority of which have been discussed in a comprehensive review by Baker *et al.* (2014)^22^. Briefly, existing bladder models are produced using one of three broad culture techniques: (1) organ culture of intact biopsies or explant culture; (2) culture of urothelial cells naturally shed into the urine or harvested from biopsies; and (3) organotypic culture whereby normal urothelial cells are stimulated to form 3D organoids on filter inserts^22^. Although arguably the most relevant model system, organ culture of intact human tissue is time consuming, yields a finite amount of experimental material and requires fresh human tissue^22,40^. Due to these limitations, a number of researchers have used rodent bladder biopsies. However, inter-species differences imply that these models may not be as biologically germane^41^. More practical is the cultivation of human urothelial cells isolated from host urine or biopsies which, when grown using a specialist protocol, have been shown to maintain the ability to stratify, differentiate and develop a robust barrier function *in vitro*^22,42-45^.

Although these models constitute impressive alternatives to animal models, to our knowledge, they last only a few hours up to a day in urine *in vitro*^22,46^, the natural apical substrate of this tissue. Therefore, the effect of urine exposure on urothelial differentiation and GAG expression remains unclear. Moreover, none of the human-derived urothelial biomimetics have been reported to correctly express cytokeratin 20 at the apical surface^35^. To address these limitations, we worked to develop a urine-tolerant organotypic human urothelium that could be used as a platform studying for host/uropathogen interactions, treatment, and resolution in humans.

## Results

### HBLAK and HBEP can form three-dimensional urothelial organoids

HBEP cells were derived from normal human bladder biopsies by CellNTec and provided commercially in cryovials. We also grew HBLAK cells, which are spontaneously immortalised but not transformed version of HBEP available from the same company. These latter cells retain the ability to differentiate but have increased longevity without senescing. On thawing, both cell types were seeded on plastic in fully defined, serum-free, BPE containing CNT-Prime media, which favours the proliferative phenotype. Both HBEP and HBLAK shared a ‘spindle-like’ morphology, a hallmark of multipotent epithelial progenitors. When 70% confluent, the cells were transferred to Millicell transwells (Millipore) with CNT-Prime media in the apical and basal chambers and grown to confluency. At this point, the media was shifted to high-calcium, differentiation media (CNT-Prime-3D) in both chambers. After overnight incubation, we replaced filter-sterilized human urine in the apical chamber and left the cultures to develop for 14 (HBLAK) to 25 (HBEP) days depending on the experiment, with periodic media and urine changes. At endpoint, the organoid-coated filters were retrieved, fixed, and stained for various biomarkers and inspected microscopically.

3D confocal analyses showed that both cultures were viable despite the prolonged presence of urine. The HBEP tissue contained multiple layers (approximately 3) with tightly-packed spheroid basal cells, intermediate cells and enlarged and flattened umbrella-like cells at the apical surface (Fig. 1a,b). The HBLAK tissue was morphologically similar, but in contrast, likely due to increased rate of growth, these produced approximately 5-7 cell layers with multiple layers of intermediate cells (Fig. 1c,d). Optical slices at the basal region of the HBEP and HBLAK organoids respectively showed the typically small, tightly-packed and spheroid morphology of urothelial basal cells (Fig. 1e,g). Basal cells in the HBLAK organoid (~ø10μm) appeared to be slightly smaller than those in the HBEP culture (~ø20μm) (Fig. 1e,g). Similarly, single optical slices at the apical regions of the HBEP and HBLAK organoids showed the formation of a large, flat and often hexagonal cellular morphology typical of well-differentiated umbrella cells (Fig. 1f and 1i for HBEP, Fig. 1h and 1j for HBLAK). As shown in Figure 1j, lower-magnification views of HBLAK show a more heterogeneous differentiation pattern, with cells elaborating distinct multi-layered zones of organoid formation with large umbrella-like cells at their surface, flanked by regions of very much smaller undifferentiated basal cell-like monolayers and occasional areas of hypertrophy.

**Figure 1.**
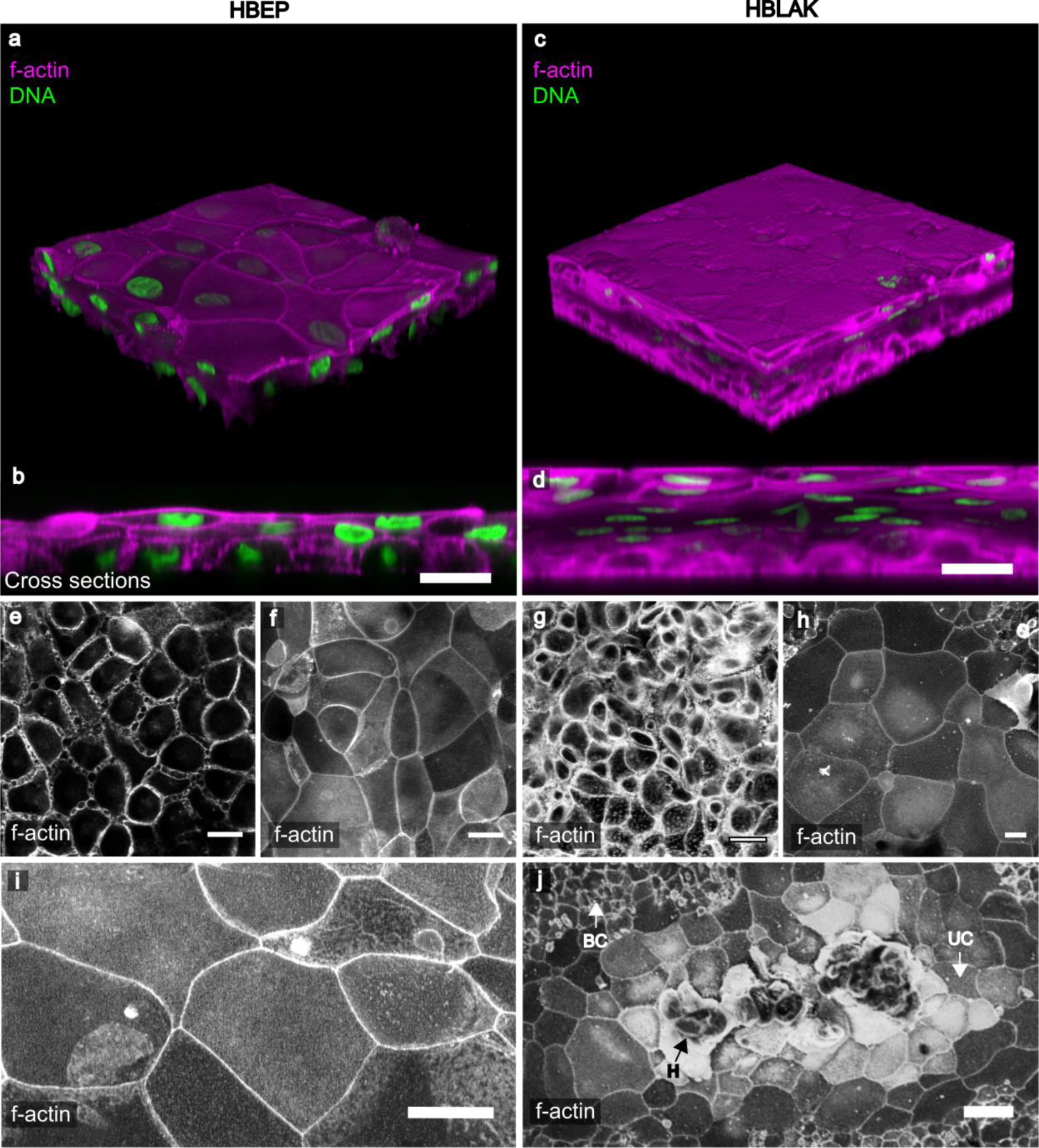
HBLAK cells retain the ability to differentiate into 3D urothelial organoids. Colour images are composites showing the phalloidin-stained F-actin in magenta and DAPI-stained DNA in green. Lower images show phalloidin-stained F-actin in monochrome. (a) 3D confocal model constructed from a 200 slice Z-stack of HBEP organoid. Umbrella-like cells are large and flat. (b) Orthogonal view of the Z-stack shows the tissue to be ~3 layers in depth and the basal cells to be spheroid in morphology. (c) 3D confocal model constructed from a ~300 slice Z-stack of HBLAK organoid. The HBLAK tissue is significantly better developed than the HBEP tissue in terms of thickness and number of cell layers. (d) Orthogonal reslice of the HBLAK tissue shows ~5-7 cell layers with flattened apical cells and more spheroid cells beneath. (e) Single optical slice at lowest region of the HBEP organoid showing small tightly packed basal cells. (f) Single optical slice at apical region of HBEP organoid showing well-differentiated, characteristically large umbrella-like cells. (g, h) Single optical slices of basal and umbrella-like cells respectively in HBLAK bladder bio-mimetic. Scale bars represent 20μm. (i) A further single optical slice showing large umbrella-like cells at the surface of the HBEP organoid. Scale bar represents 20μm. (j) Maximum projection confocal image of a region of HBLAK organoid formation. Large, flat umbrella-like cells (UC) lined the apical surface of the organoid surrounded by monolayers of small undifferentiated basal-like cells (BC). Areas of hyperplasia (H) were noted at the ‘peak’ of a small proportion of investigated regions of differentiation. Scale bar represents 40μm.

In summary, both models formed large zones of three-dimensional epithelia in a manner reminiscent of a urothelium, with umbrella-like cells at their apical surface. Moreover, both models were exposed to sterile human urine for several weeks (14-25 days) without exhibiting any signs of toxicity.

### HBLAK and HBEP organoids exhibit key biomarkers of the human urothelium

We characterised the HBEP and HBLAK human urothelial organoids further by targeting urinary tract-specific antigen expression using indirect IF in conjunction with high-resolution laser scanning confocal microscopy. The resulting organoids had the correct spatial expression of several key biomarkers. Studies exploring cytokeratin (CK) expression in normal human bladders have shown a relationship between the level of cytodifferentiation and sub-type CK expression^18^. Both HBEP‐ and HBLAK organoids exhibited correct spatial expression of CK8 and CK20; specifically, CK8 was expressed throughout the strata of the *in vitro* tissues whereas CK20 was expressed preferentially by the umbrella-like cells at the apical surface (Fig. 2 a,b,c,d). In contrast to the rodent bladder, muscarinic receptors are expressed throughout the human urothelium^47,48^. This finding was echoed in our model with evidence of muscarinic receptors M2 and M3 expressed in all cell layers of both organoids (Fig. 2e and g). Uroplakin-III (UP3), an indispensible part of the asymmetric unit membrane^49^, was present at the apical surface of the HBEP (Fig. 2f) and HBLAK (Fig. 2h) organoids, in a speckled pattern (HBEP, Fig. 2i; HBLAK, Fig. 2k) similar to that seen in urothelial cells shed in the urine of a chronic UTI patient (Fig. 2j,l). Multinucleation of the umbrella-like cells was a relatively rare occurrence (e.g. examples can be seen in Fig 2g, i and k).

**Figure 2.**
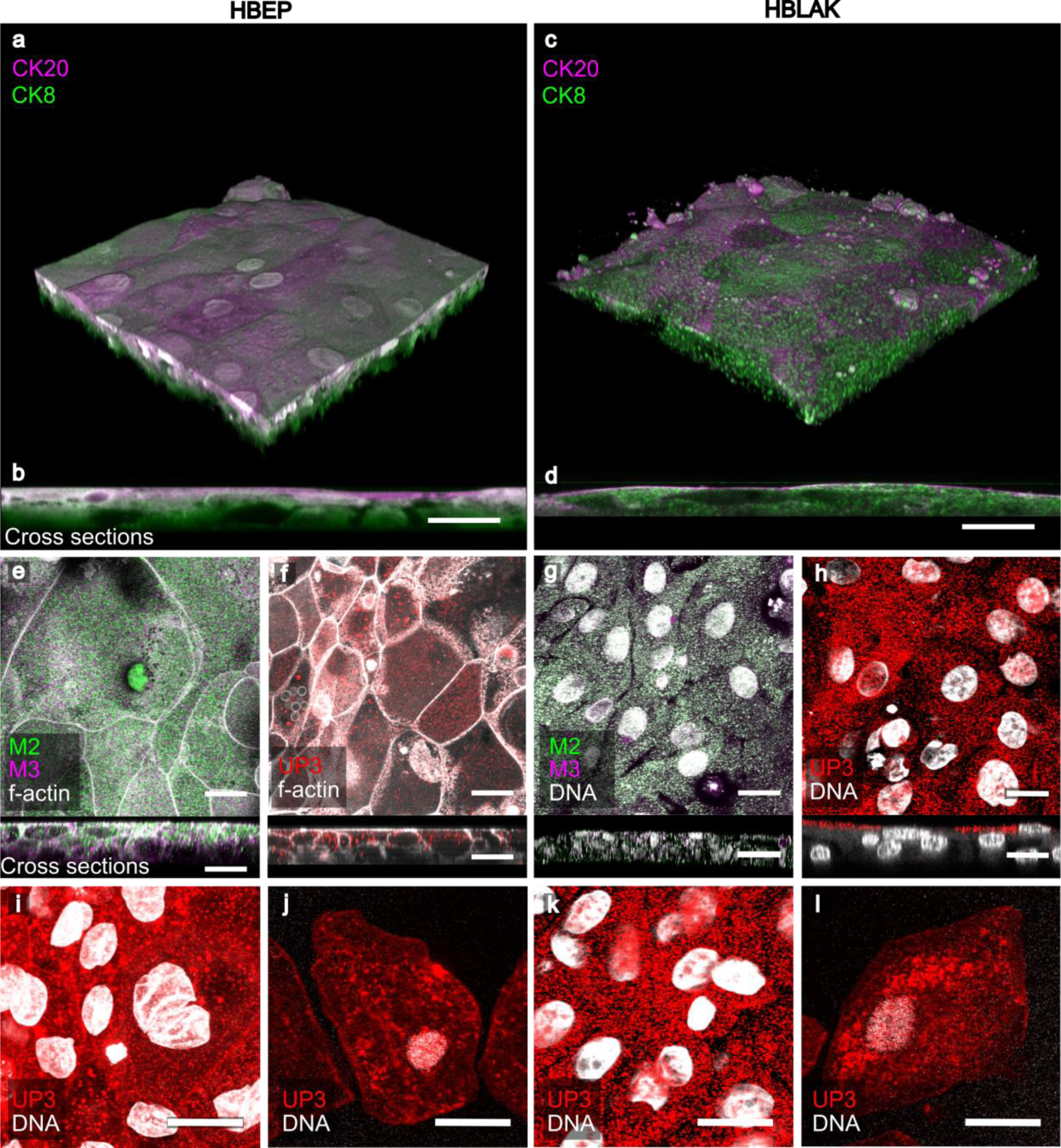
Characterisation of HBEP (left) and HBLAK (right) urothelial organoids using IF. (a) 3D confocal model and (b) orthogonal reslice from a 100 slice Z-stack of HBEP organoid. The HBEP organoid exhibited the correct spatial expression of Cytokeratin-20 (CK20, umbrella cells, magenta) and Cytokeratin-8 (CK8, throughout urothelium, green). (c) 3D confocal model and (d) orthogonal reslice from a 120 slice Z-stack of HBLAK organoid. As with the primary HBEP mimetic, the HBLAK organoid also expressed CK20 (magenta) at the apical surface and CK8 (green) throughout the tissue. (e-h) Single optical slices at apical region of HBEP (e,f) and HBLAK (g,h) organoids with corresponding orthogonal cross sections shown directly beneath. (e,g) Expression of muscarinic receptor 2 (M2, green) and muscarinic receptor 3 (M3, magenta). Both receptors were found throughout the tissue in both models. Phalloidin-stained F-actin is presented in grey. (f,h) Expression of Uroplakin-III (UP3, red) preferentially at apical region of both models. Phalloidin-stained F-actin is presented in grey. (i) High-power single optical slice of UP3 (red) expression in HBEP model. (j) Pattern of UP3 (red) expression found in an exfoliated urothelial cell harvested from a chronic UTI patient. (k) High-power single optical slice of UP3 (red) expression in HBLAK model. (l) Further example of the pattern of UP3 (red) expression found in patient-isolated urothelial cells. DAPI-stained DNA is presented in grey. Scale bars represent 20μm.

### HBLAK organoids possess correct topographical and ultrastructural features

Given these promising results, we went forward with the more tractable and thicker HBLAK urothelial mimetic model, using scanning electron microscopy (SEM) and transmission electron microscopy (TEM) to analyse its ultrastructure, organisation and overall topography. Low-power SEM echoed what was seen in Figure 1j, namely that the HBLAK cells elaborated distinct multi-layered zones of organoid formation with umbrella-like cells at their surface (approximately two-thirds of the tissue, 64.8% +/− 10.4) along with zones of undifferentiated basal cell-like monolayers (Fig. 3a) and areas of disorganised hyperplasia (Fig. 3b). The surface of each differentiated zone exhibited very large (up to ~80μm), often hexagonal umbrella-like cells (Fig. 3c,d,e). A comparison with the undifferentiated basal cells (Fig. 3e vs. f) showed that the umbrella-like cells were up to approximately 50 times larger in terms of surface area than their undifferentiated counterparts. SEM micrographs also showed evidence of tight junction formation (Fig. 3c)^50^ and structures resembling, in size and spacing, characteristic microplicae or ‘hinges’ at the apical surface of each umbrella-like cell (Fig. 3g)^50,51^. Orthogonal sections of fully differentiated organoid zones were analysed using TEM. As with the laser scanning confocal imaging, TEM elucidated distinct layers of basal, intermediate and umbrella cells (Fig. 3h). Structures consistent in size and spacing with rigid plaques could be seen residing between the ‘hinges’ at the apical surface of the umbrella cells (Fig. 3i,j)^50,51^. Structures were also seen that may correspond to the specialised fusiform vesicles (Fig. 3j) necessary for trafficking uroplakins^52^. In contrast, TEM analysis of the small, undifferentiated cells making up the monolayers flanking the organoid zones did not exhibit the putative hinges, plaques or fusiform vesicles (Fig. 3k,l). Taken together, these observations suggest that the fully differentiated regions of the organoids possess some key features expected of a human urothelium.

**Figure 3.**
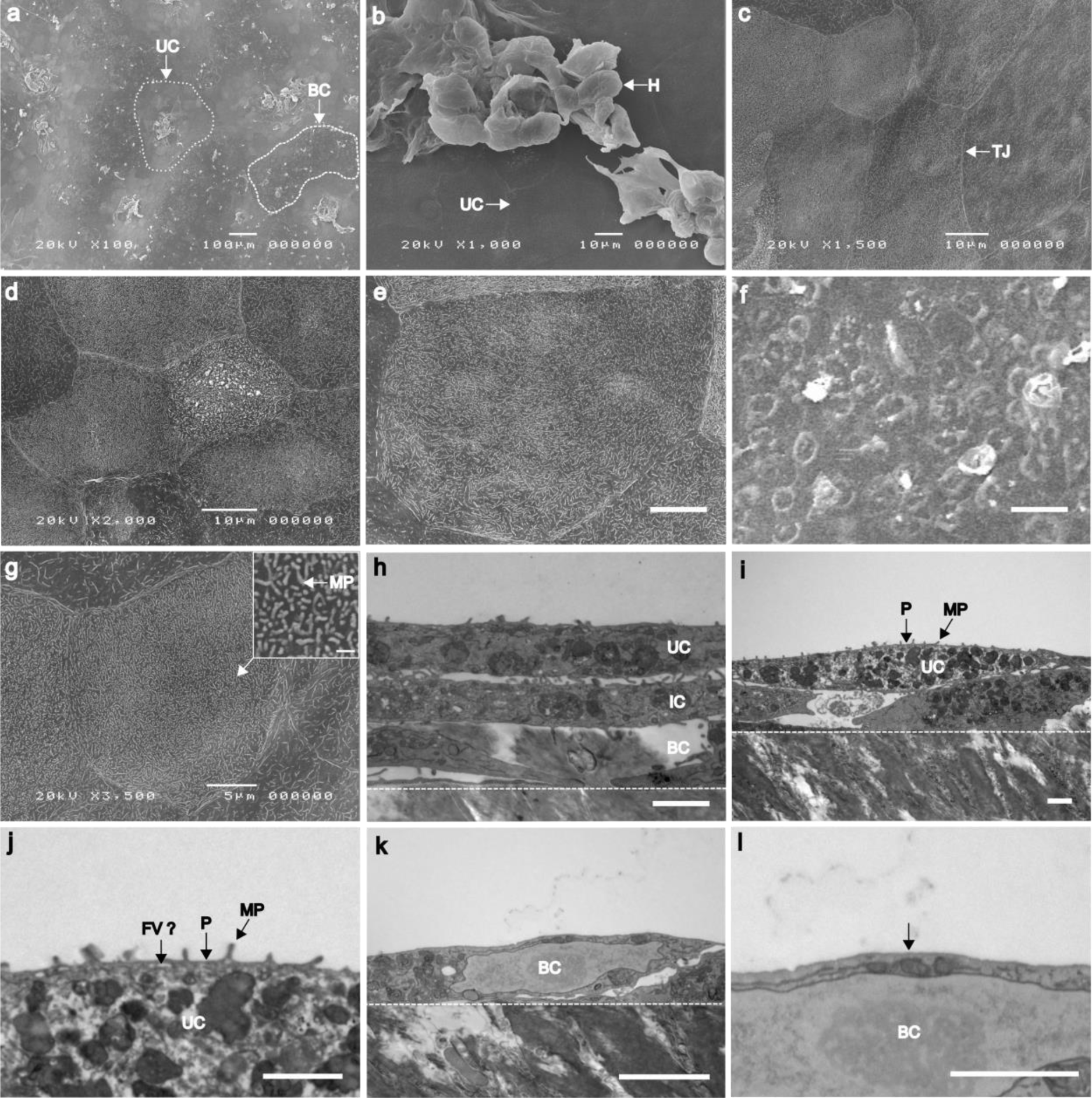
Analysis of HBLAK organoid topography and ultrastructure using scanning electron microscopy (SEM) and transmission electron microscopy (TEM) (a) SEM micrograph showing ‘islands’ of organoid formation. Umbrella-like cells (UC) can be seen at the apical surface of each ‘island’ interspersed by areas of undifferentiated monolayers of basal-like cells (BC). (b) SEM at the apical surface of organoid. Large (~50-60μm), flat umbrella-like cells (UC) are present at the upper-most surface of each ‘island'. Areas of hyperplasia (H), however, were frequently observed. (c) SEM of umbrella-like cells showing the formation of tight junctions (TJ). (d-e) SEM of further regions of large, characteristically tessellated umbrella-like cells. Scale bars represents 10μm. (f) SEM of region of small undifferentiated basal-like cells. Scale bar represents 10μm. (g) SEM of a single umbrella cell. Microplicae (MP) or ‘hinges’ can be seen covering the surface of the umbrella-like cells. Inset scale bar represents 500nM. (h) TEM of 3-4 layered organoid. Basal cells (BC) can be seen above the polycarbonate culture filter (white broken line). Large, flat intermediate cells (IC) and umbrella-like cells (UC) are superior to the basal cells. Scale bar represents 2μm. (i) TEM of organoid. Plaques (P) are present between the microplicae (MP) of the umbrella-like cell (UC). White broken line represents the upper surface of the polycarbonate culture filter. Scale bar represents 2μm. (j) Enlarged version of TEM image j. Rigid plaques (P) are situated between each hinge (microplicae, MP). White structures were evident that might be fusiform vesicles (FV?), responsible for trafficking uroplakin to the umbrella cell plaques. Scale bar represents 1μm. (k) TEM of a monolayer of undifferentiated basal cells (BC, as highlighted in image a) found between the organoid ‘islands'. These non-differentiated cells measure ~8μm across, making them approximately 10-50 times smaller than the umbrella-like cells in the same system. White broken line represents the upper surface of the polycarbonate culture filter. Scale bar represents 2μm. (l) TEM of the apical surface of the basal cell (BC) shown in image i. Microplicae, plaques and potential fusiform vesicles were not seen (white arrow). Scale bar represents 1μm.

### Urine strongly influences HBLAK differentiation, organoid development and GAG formation

As shown above, the HBLAK organoid expresses key urothelial markers and is morphologically reminiscent of a urothelium. Of key importance, however, is the tolerance exhibited by these cultures to urine. In this group of experiments we investigated whether urine is merely tolerated or indeed necessary for differentiation in 2D, organoid formation and the elaboration of a GAG layer. To achieve this, we exposed the cells, in 2D and 3D, to various dilutions of sterile human urine before examining microscopically.

To decouple the influence of urine from any effects that might be exerted by developing intermediate cells below, we grew HBLAK cells as 2D monolayers on chamber slides and exposed them to medium containing urine. As shown in Figure 4, HBLAK cells exhibited a marked dose response to urine after 72 hours. Cells grown in high calcium medium alone took on a small, tightly packed basal cell-like appearance with visually well-formed intercellular junctions as assessed by actin staining (Fig. 4a) and the elaboration of actin-containing microvilli (Fig. 4a inset, red arrow). With the addition of 25% human urine, however, a small proportion of the cells began to exhibit an enlarged morphology, but intercellular junctions were partially disrupted (Fig. 4b). After 72 hours in 50% sterile human urine, extensive colonies of HBLAK cells took on a large, flat umbrella cell-like morphology (Fig. 4c) devoid of microvilli (Fig 4c inset, red arrow). However, intercellular junctions seemed almost entirely compromised, possibly because increased urine came at the expense of calcium concentration.

**Figure 4.**
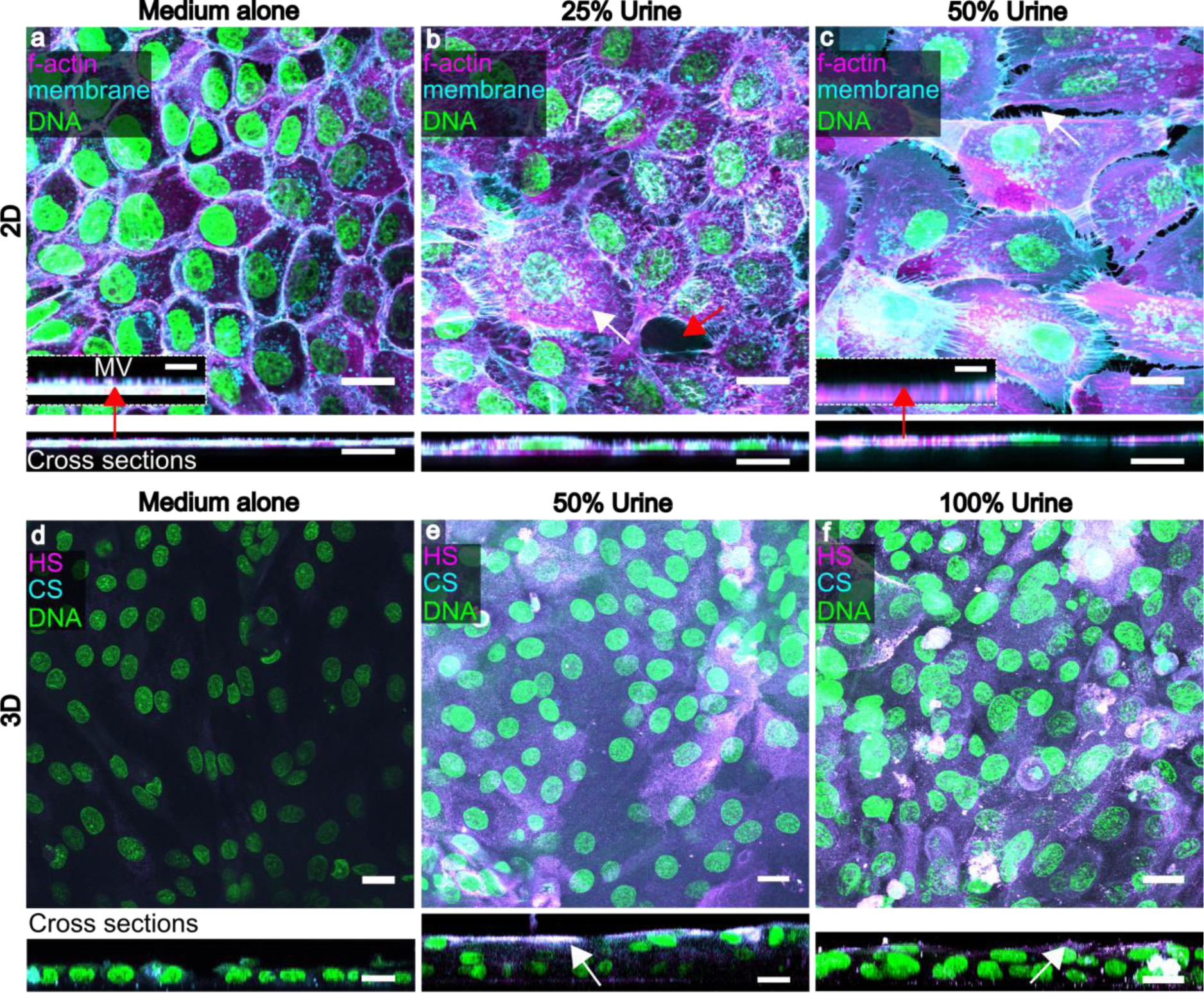
Human urine affects HBLAK differentiation, 3D organoid formation and GAG expression. Upper images are maximum projections and corresponding orthogonal cross sections (below) from 12 slice Z-stacks of HBLAK cells cultured in 2D on glass. Cells were grown to confluency and incubated for 72hrs in varying proportions of human urine in culture medium. WGA-stained plasma membrane is presented in cyan, phalloidin-treated F-actin in magenta and DAPI-labelled DNA in green. (a) Monolayer grown in 3D barrier medium alone. Basal cell morphology was maintained and intercellular junctions appeared to be intact. Inset displays high magnification cross section showing actin-rich microvilli (MV) at apical surface. Inset scale bars represents 5μm. (b) HBLAK monolayer cultured in 25% human urine. A subset of cells began to differentiate (white arrow) exhibiting a large, flat umbrella cell-like morhology. Cell junction integrity was disrupted (red arrow). (c) HBLAK monolayer cultured in 50% human urine. Large colonies of cells exhibited an umbrella cell-like morphology. Cell junctions, however, were almost entirely compromised (white arrow). Inset shows the lack of microvilli on the surface of the umbrella-like cells. Inset scale bars represents 5μm. Lower images are maximum projections (and cross sections below) from 20-slice Z-stacks of HBLAK cells grown on 3D culture filter inserts. Cultures were exposed to 3D barrier medium in the basal compartment and varying proportions of human urine at their apical surface for 14 days. The glycosaminoglycan (GAG) constituents heparan sulphate (HS) and chondroitin sulphate (CS) were labelled and are shown here in magenta and cyan respectively. DAPI-stained DNA is shown in green. (d) Cells cultured in 3D barrier medium only in the basal and apical compartments. No stratified organoid formation was observed. Little GAG expression was seen. (e) Cells cultured with 50% urine at the apical surface. Urothelium-like organoids were formed. Heperan sulphate and chondroitin sulphate were strongly expressed by umbrella-like cells (white arrow). White represents colocalization. (f) Cells cultured with 100% urine at the apical surface. Again, well organised bladder organoids were formed and a GAG (heparan sulphate and chondroitin sulphate) mucin layer was elaborated at the cell-urine interface (white arrow). Scale bars represent 20μm.

Strikingly, we found human urine to be necessary for organoid formation and the elaboration of a GAG layer in HBLAK cells. Cells cultured for 14 days on filter inserts with high calcium medium in the basal and apical compartments were comprised of only one layer, showing no stratified organoid formation, little heparan sulphate, and no detectable expression of chondroitin sulphate (Fig. 4d). Moreover, these cells appeared to exhibit a degree of anaplasia, with a more ‘spindle-like’ multipotent progenitor cell-like morphology (data not shown). In contrast, HBLAK cells cultured in 50% or 100% human urine at the apical surface produced zones of organoid formation between 3 and 6 cell layers thick (Fig. 4 e,f) as previously seen in the experiments presented in Figure 1, and both stimulated the expression of a heparan and chondroitin sulphate-rich GAG layer at the urine-umbrella cell interface (Fig. 4 e,f). Taken together with the data in Figure 3, these experiments suggest that urine is an indispensable effector of urothelial differentiation in this model. Nevertheless, despite a lack of GAG layer and of the structures that appear to be asymmetric unit membrane plaques, undifferentiated cells remain viable for long periods in urine, which suggests that these “barriers” are not required for urine tolerance.

### The HBLAK organoid is a promising model for studying *Enterococcus* infection

Our laboratory studies the uropathogen *Enterococcus faecalis*. This bacterium is commonly implicated in chronic UTI in the elderly and is frequently associated with multi-drug resistance, hospital-acquired infection, and catheter-associated biofilm formation; we previously demonstrated that it exhibits intracellular invasion in patient cells^53-61^. To determine whether the organoid is a good model for studying host-pathogen interactions with this bacteria, we infected the HBLAK organoid with patient-isolated *E. faecalis*^54^. When we inspected the cultures two hours after infection, we found that *E. faecalis* exhibited robust tropism in this model, with the resulting adherent colonies relatively loosely packed (Fig. 5a). As with human patients suffering from acute and chronic UTI, the apical cell layer was shed in response to bacterial insult^54^, leaving an uneven surface of basal and intermediate cells (Fig. 5a). Inspection of supernatants post-infection revealed extensive shed umbrella-like cells (Fig 5b) reminiscent of those seen in the urine of infected patients (Fig. 5c and^54^). Significantly, in the tissue that remained, we saw frequent examples of large intracellular bacterial colonies (Fig. 5d, arrows) within the superficial layer of cells, showing a similar loosely-packed morphology to those we previously observed in shed urinary umbrella cells from patients^54^. Image analysis showed that 7.3% of cells harboured intracellular colonies with a mean of 37.8 (+/− SD of 11.5) bacteria per cell. A gentamicin protection assay performed on organoids infected with *E. faecalis* for two hours (Fig. 5e) supported this observation, with viable intracellular bacteria liberated after extracellular sterilization and host cell lysis. Specifically, treatment with 2000μg/ml of gentamicin resulted in zero detected growth within the organoid supernatant. Detergent treatment liberated intracellular bacteria resulting in the detection of a median of 4.8×10^3^ cfu/ml in the lysate (approx. 55% bacterial recovery). A non-parametric Friedman test of differences among repeated measures was conducted on the median cfu/ml data from the untreated (control, N=3) and gentamicin treated (N=3) organoids. A statistically significant difference was found among the gentamicin treated organoids (χ^2^=6, p=.04) whereas no difference was found among the untreated control data (χ^2^=3.8, p=.15). Taken together, these findings suggest that the model recapitulates some key aspects the host/pathogen interaction of *E. faecalis* with its human host.

**Figure 5.**
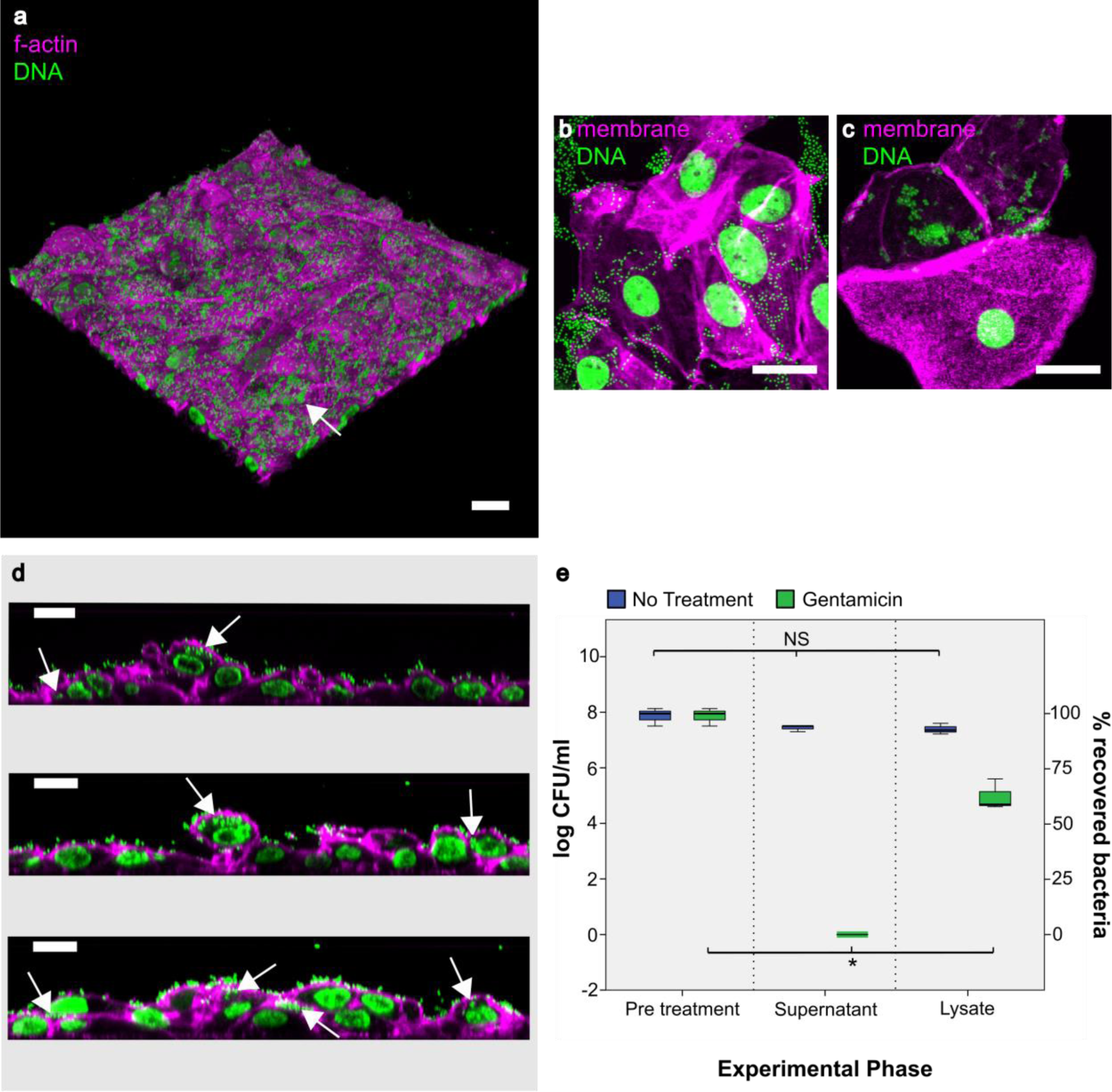
Analysis of HBLAK-derived urothelial organoids infected with uropathogenic *E. faecalis*. Composites showing (a) 3D confocal model constructed from a 100 slice Z-stack of HBLAK organoid post infection with *E. faecalis* (MOI of 25). phalloidin-stained F-actin in magenta and DAPI-stained DNA (host and pathogen) in green. The characteristically smooth and flat umbrella-like cell layer was lost in response to infection. Loose colonies of *E. faecalis* could be seen adhering to the now unprotected basal and intermediate cells (white arrow). (b) Surface cells shed from organoid in response to infection with *E. faecalis.* WGA-labelled membrane shown in magenta and DAPI-stained DNA (host and pathogen) in green. (c) Patient-isolated urothelial cells shed into the urine in response to infection. (d) Orthogonal cross sections prepared from image a. White arrows show evidence of intracellular *E. faecalis* within the basal and intermediate cells of the HBLAK organoid. 7.3% of cells exhibited intracellular colonisation with a mean of 37.8 (+/− SD of 11.5) bacteria per cell. Phalloidin-stained F-actin in magenta and DAPI-stained DNA (host and pathogen) in green. Scale bars represent 20μm. (d) Box and whisker plot showing the results of gentamicin protection assays undertaken on HBLAK organoids infected with uropathogenic *E. faecalis*. Results are represented as log10 median CFU/ml with a % bacterial recovery derived axis. Treatment with 2000μg/ml of gentamicin resulted in zero detected growth within the organoid supernatant. Detergent treatment liberated intracellular bacteria resulting in the detection of 4.8×10^3^ cfu/ml in the lysate (approx. 55% bacterial recovery). N=3 per treatment. NS = No statistically significant difference (P>.05), *P≤.05.

In summary, this model, to our knowledge, represents the first long-term urine-tolerant human bladder organoid produced *in vitro*. This tissue is reminiscent of normal human bladder urothelium and expresses a number of key markers in the correct spatial compartments in response to exposure to urine. As with patients, this organoid rapidly sheds the apical cell layer in response to bacterial insult. Moreover, *E. faecalis* displayed invasive phenotypes, supporting our previous findings in shed patient cells^54^.

## Discussion

The development and use of *in vitro* human tissue mimetics is thought to be accelerating the drug discovery process^62^ and improving our understanding of human tissue morphogenesis. Due to improved physiological relevance, such models could even reduce the use of animal models in the coming years^63^. Here, we present a urine-tolerant, three-dimensional urothelial organoid derived from human progenitors that is easy to grow from commercially-sourced, quality-controlled materials, that displays key hallmarks of the human urothelium, and which may serve as an alternative to the murine model. If desired, it should also be possible to create similar urine-tolerant organoids from fresh human biopsies using the protocols and defined medium we describe. An even more physiological response could well be achieved by embedding the organoid into a perfusion-based bioreactor allowing the flow of apical urine, which would introduce relevant shear forces and metabolite flux^46^.

As this organoid model was generated in a similar manner to others that have been reported^22^, it is not clear why it has superior urine tolerance. Additional work in our laboratory has not revealed any differences in outcome depending on the source of the human urine. The primary cells, and their HBLAK derivatives, were obtained from standard healthy adult human biopsy of the trigome region, and we do not anticipate this being materially different from other biopsies; indeed, we achieved good results with similar cells sourced from a different company. Therefore, the most likely factor is the specialized medium, which is free of serum and other growth factor supplements that could impair differentiation. We are currently using an empirical approach to determine which aspects of the growth media and urine are important for differentiation.

Morphologically, the HBEP and HBLAK cells produced tissue with encouraging similarities to human urothelium^18,19,22,26^. HBEP cells produced tissue ~3 cell layers thick whereas HBEP cell-derived tissue elaborated ~5-7 layers. In contrast to mice, higher mammals such as humans have multiple intermediate cell layers, a feature which may favor the use of HBLAK cells^21^. However, rate of cell division could have played a role in this outcome, as could cell senescence in the HBEP population^64^. Both culture types developed an apical layer of enlarged, flattened cells that appeared umbrella-cell-like^21^ in morphology. Multiple nuclei were not very common in these cells, as would be expected from the mouse urothelium. We could find no consensus in the literature about the expected percentage of multinucleation in the human urothelium, with studies being scant, and it is also not clear whether the multinucleated state plays any functional role. Further work is needed to understand how tissue thickness and multinucleation is regulated in both cell types.

Phenotypic analysis of both the HBEP and HBLAK urothelial organoid tissue elucidated the presence of some important urothelial markers. Uroplakins are a group of highly conserved glycoproteins which are unique to mammalian urothelium^49^. Uroplakin-III (UP3) was expressed on the apical surface throughout the differentiated tissue and its expression was morphologically similar to that of patient-shed urothelial cells. Laguna *et al.* (2006)^18^ found that the normal human urothelium expresses a range of cytokeratins in relation to the level of cytodifferentiation. Our 3D culture mimicked these findings with cytokeratin-8 (CK8) found throughout the tissue and cytokeratin-20 (CK20) being a preferential phenotype of well-developed umbrella cells^18^. To our knowledge, this is the first human bladder organoid demonstrating the correct spatial expression of CK20^35^.

Patients with overactive bladder symptoms are frequently treated with antimuscarinics which target muscarinic receptors in the detrusor and urothelium^47^. Crucially, rodent bladder urothelial cells do not express muscarinic receptors^48^. In the case of the HBEP and HBLAK tissue grown in this study, muscarinic receptors M2 and M3 were detected throughout the urothelial cell layers, further supporting its physiological relevance and potential use in the development of novel therapeutic agents for the bladder symptoms of MS and other neurogenic disorders.

More detailed analysis revealed several interesting aspects to our HBLAK organoid model. First, EM showed that the tissue was not homogenously differentiated; instead, the surface consisted of three discrete zones: ‘valleys’ of undifferentiated monolayer; ‘plateaus’ of fully differentiated 3D tissue; and ‘mountains’ of hyperplasia. Second, inspection of GAG layer markers revealed that the presence of urine correlates with its elaboration, in parallel with differentiation and organoid formation. Third, focusing on the differentiated plateaus, which comprised about two-thirds of the total area, our EM imaging shows expected key features, including morphologically distinct layers, and the enlarged flat nature of the distended umbrella-like cells. We also noted structures consistent in shape, size and spacing with the characteristic hinges and plaques associated with the AUM, and with the fusiform vesicles responsible for trafficking during bladder filling and emptying phases. These structures were entirely absent from the zone of undifferentiated cells, showing a possible correlation with differentiation status. However, higher resolution EM imaging and immunostaining is needed to confirm their identity.

Intriguingly, when we tested the barrier function of the organoid by several methods, including transepithelial resistance and fluorescent dextran permeability (data not shown), we found a lack of what is traditionally thought of as urothelial “barrier function”. We presume that this result was caused by the sporadic presence of undifferentiated cells across the tissue, which in essence short-circuit the apicobasal electrical potential difference. This result, taken together with the fact that the non-differentiated zone devoid of AUM-like structures and GAG can grow for several weeks in the presence of urine, strongly suggests that AUM and GAG are not required for urine tolerance, and that something intrinsic in the cells themselves confer resistance to its toxic effects. It also suggests that apical urine is necessary for full differentiation. Whilst it may be the case that urine is not sufficient for differentiation, given the presence of the undifferentiated ‘valleys’, it may also be the case that a subset of HBLAK cells are subtly different. Further exploration into the nature of urine tolerance and the effect of urine and other media components on differentiation and hyperplasia are warranted. Genomic analysis of cells from various regions, compared with starting cells, would also be interesting. In the meantime, researchers performing image-based studies should be able to focus on areas of the organoid corresponding to their desired level of differentiation.

Our model shows great promise for studying the host/pathogen interactions of UTI in a human-cell system. The majority of host/pathogen interaction studies in UTI focus on the most common uropathogen *E. coli*, but very little is known about other uropathogens such as *Enterococcus faecalis*, which is more common in certain cohorts, such as the elderly, the hospitalized and those using urinary catheters. We previously reported the discovery of intracellular colonies of *E. faecalis* harbored within the urothelial cells of chronic UTI patients^54^. This previous report also slowed sloughing of umbrella cells from the epithelial lining into the urine, which is known in both mice and humans to be a common response to infection^65-69^. These findings were echoed in our urothelial organoid, where *E. faecalis* formed significant intracellular colonies within the intermediate and basal cells of the urothelial mimetic after its umbrella-cell layer had been jettisoned. These results further support the notion that *E. faecalis* exhibits an intracellular phenotype.

In conclusion, current advances in 3D tissue culture enabled us to grow physiologically relevant, organotypic human models of the bladder. Human bladder biomimetics could be used as a reproducible test bed for chronic infective disease formation, treatment, and resolution in humans.

## Materials and methods

### Human Primary progenitor cell expansion and handling in 2D

Commercially available human bladder epithelial progenitor cells (HBEP, Cell N Tec)^70^ and their spontaneously immortalised, non-transformed counterparts (HBLAK, Cell N Tec) were supplied in frozen aliquots containing ~5×10^5^ cells at passage 2 and ~0.5×10^5^ at passage 25 respectively. Cells were isolated from bladder trigone biopsies from male patients undergoing surgery for benign prostatic hyperplasia. HBEP cells are guaranteed to grow for a further 15 population doublings before senescing whereas HBLAK cells, although spontaneously immortalised, should not be differentiated into 3D cultures once they have exceeded a passage number of 40-50. Both cell types were cultured identically, with the exception of slight differences in incubation time between passages, due to the slightly increased rate of cell division exhibited by HBLAK cells.

Thawed cells were seeded (~300 cell clumps / cm^2^) into pre-warmed and equilibrated low-calcium, high-bovine pituitary extract, primary epithelial medium (CnT-Prime, Cell N Tec) in 9cm polystyrene dishes and incubated at 37°C in a humidified incubator under 5% CO_2_. Culture medium was replaced after overnight incubation to remove residual dimethyl sulfoxide (DMSO). Antibiotics were not added to culture medium at any point due to adverse effects on cytodifferentiation, metabolism and morphology ^71^. Furthermore, trypsin is known to damage primary cells, so Accutase solution (Innovative Cell Technologies) was used to detach cells at all stages of experimentation^72^. Cells were allowed to expand to ~70% confluency before freezing batches of cells at a density of ~1×10^6^ cells/ml in defined freezing medium (CnT-CRYO-50, Cell N Tec) in preparation for later experiments. Cells were not allowed to become fully confluent during cell expansion in an effort to maintain a proliferative phenotype.

### Differentiation of 3D human urothelium *in vitro*

In preparation for organotypic culture, previously frozen progenitor cells were thawed and expanded on 9cm culture dishes as above. Once 70-80% confluent, the cells were washed briefly with calcium‐ and magnesium-free phosphate buffered saline (PBS, Sigma-Aldrich) and incubated at 37°C in ~3ml of pre-warmed Accutase solution for 2-5min. The dishes were lightly tapped and detached cells re-suspended in 7ml of warm CnT-Prime. After centrifugation at 200xg for 5min, the supernatant was removed and the pellet re-suspended in fresh CnT-Prime. This cell suspension was counted whilst allowing the cells to equilibrate for 3min at room temperature. 2×10^5^ cells in 400μl of CnT-Prime (internal medium) were added to 6 12mm 0.4μm pore polycarbonate filter (PCF) inserts (Millipore) standing in 6cm culture dishes containing ~3ml of fresh pre-warmed CnT-Prime medium (external medium, level with insert filters). A further 8ml of CnT-Prime medium was added to the 6cm dish (external to the filter inserts) until internal and external fluid levels were the same. The 3D culture inserts were incubated for 3-5 days until 100% confluent. Confluency was determined through the fluorescent staining of 1 insert and visualisation under epi-fluorescence microscopy (see section below). Once deemed confluent, internal and external medium was removed and replaced with low-BPE, calcium-rich (1.2mM) differentiation barrier medium (CnT-Prime-3D, Cell N Tec) to promote differentiation. Subsequent to overnight incubation, the internal medium (apical surface of cell culture) was removed and replaced with filter-sterilised human urine pooled from healthy volunteers of both genders to aid terminal differentiation into umbrella cells. The external CnT-Prime-3D medium and the internal human urine were replaced every 3 days and the culture incubated for 14-24 days at 37°C in 5% CO_2_.

To explore the effect of urine on differentiation in 2D, the HBLAK cells were seeded on 8-well permanox Lab-Tek slides (Sigma-Aldrich) and grown to confluency. The cells were then exposed to CnT-Prime-3D medium alone, 25%, or 50% sterile human urine diluted in CnT-Prime-3D for 72 hours at 37°C in 5% CO_2_. To analyse the effect of urine on HBLAK organoid formation, cells were grown to confluency on filter inserts as above. The basal compartment was treated with CnT-Prime-3D throughout, however, the apical compartment was filled with either CnT-Prime-3D alone, 50% sterile human urine diluted in CnT-Prime-3D or 100% urine. The specified medium or urine was changed every 3 days and the culture incubated for 14 days at 37°C in 5% CO_2_.

### Characterisation of the 3D urothelium

Prior to fluorescent staining and immunofluorescence (IF), filter inserts were carefully transferred to 8-well plates (Nunc) and submerged in 4% methanol-free formaldehyde (Thermo Scientific, Fisher Scientific) in PBS overnight at 4°C. After fixation, the filter inserts were kept at 4°C in 1% formaldehyde in sealed containers in preparation for processing.

To determine confluency and analyse morphology, the pre-fixed tissue was permeabilised in 0.2% Triton-X100 (Sigma-Aldrich) in PBS for 15 minutes at RT followed by a single wash with PBS. The cells were stained with TRITC or Alexa Fluor-633-conjugated phalloidin (0.6μg/ml)(Sigma-Aldrich), to label filamentous actin, and the DNA stain 4’’,6-diamidino-2-phenylindole, (DAPI, 1μg/μl; Sigma-Aldrich) in PBS for 1 hour at RT. The dual-labelling solution was gently aspirated and the cells washed 5 times in PBS.

For indirect IF, the tissue was permeabilised as above, washed with PBS then blocked with 10% normal goat serum (NGS, Thermo Fisher) in PBS for 1 hour. Tissue was incubated overnight at 4°C with primary antibodies in PBS containing 1% NGS as follows: 1:10 dilution of mouse anti-uroplakin-III (UP3) monoclonal antibody (clone AU1, 651108, Progen Bioteknik); 1:50 dilution of mouse anti-Cytokeratin 8 (CK8) monoclonal antibody (clone H1, MA1-06317, Thermo Fisher); 1:100 dilution of rabbit anti-Cytokeratin 20 (CK20) polyclonal antibody (PA5-22125, Merck Millipore); 1:200 dilution of rat anti-muscarinic acetylcholine receptor m2 (M2) monoclonal antibody (clone M2-2-B3, Merck Millipore); 1:200 dilution of rabbit anti-muscarinic acetylcholine receptor m3 (M3) polyclonal antibody (ab126168, Abcam); 1:100 dilution of mouse anti-chondroitin sulphate monoclonal antibody (clone CS-56, ab11570, Abcam) or 1:100 dilution of rat anti-heparan sulphate proteoglycan (large) monoclonal antibody (clone A7L6, ab2501, Abcam). Post incubation with primary antibodies, the tissue was washed 5 times with PBS containing 1% NGS then incubated at RT for 1 hour with a 1:250 dilution of the following secondary antibodies (depending on the species of primary antibody used): goat anti-mouse, goat anti-rabbit or goat anti-rat conjugated to either Alexa Fluor-555, Alexa Fluor-488 or Alexa Fluor-633 (Invitrogen). Labelled cells were washed 5 times with PBS to remove unbound secondary antibody before staining with phalloidin and DAPI as above. In some experiments, prior to permeabilization, cell plasma membranes were labelled with 1μg/ml wheat germ agglutinin (WGA) conjugated to Alexa Fluor-488/633 (Invitrogen) in Hank’s balanced salt solution (HBSS, Invitrogen) for 20min at RT. Controls were performed by using primary and secondary antibodies in isolation.

In preparation for imaging, filters were carefully removed from inserts using a scalpel, mounted with FluorSave reagent (Calbiochem), and a coverslip fixed in place with clear nail varnish. Lab-Tek slide wells and gaskets were carefully removed prior to the addition of FluorSave and a coverslip as above.

### Electron microscopy

Electron microscopy was conducted by the Division of Medicine, University College London electron microscopy unit at the Royal Free Campus, Hampstead, London.

For transmission electron microscopy (TEM), samples were fixed in Karnovsky’s fixative (2.5% glutaraldehyde / 2% paraformaldehyde) and then washed in 1M PBS 3×10 min, followed by a post-fixation in 1% Osmium tetroxide for 1 hour at room temperature. Tissue was rinsed with distilled water 3×10 min. Samples were dehydrated in an ethanol series (30, 50, 70, 90 and 100%) then treated with resin-ethanol (1:1) overnight. Subsequently, samples were embedded in 100 % LEMIX resin and incubated at 65ºC for 24 hours. Ultra-thin sections of the resin block were cut and post stained with 2% Uranyl acetate and Lead citrate.

For scanning electron microscopy (SEM), the samples were fixed and dehydrated as above. Samples were then incubated in Tetramethylsilane for 10 min and air dried before mounting on stubs and sputter-coated with gold.

### Experimental infection of the human urothelial organoid and the gentamicin protection assay

A single strain of *Enterococcus faecalis (E. faecalis)* originally derived from a patient with chronic UTI^54^ was grown aerobically in a shaking incubator at 37°C for 24 hours. Once a batch of 6 HBLAK 3D urothelial cultures had reached 14 days of growth, 1.6×10^7^ colony-forming units of each bacteria (MOI of 25) were added to the filter-sterile human urine at the apical liquid-liquid interface of each culture. The experimentally infected cultures were incubated for 2 hours at 37°C under 5% CO_2_. The 3D culture filter inserts were washed with PBS before fixation and staining as above. For the gentamicin protection assay, after the infection was completed, the cultures were washed 3 times in PBS before the addition of 3D barrier medium containing 2000μg/ml of gentamicin (Sigma-Aldrich) to the apical and basal compartments of the culture filter. The organoids with antibiotics were incubated for a further 2 hours at 37°C under 5% CO_2_ to kill any extracellular bacteria. Post incubation the supernatant was serially diluted in PBS (undiluted, 1:100, 1:1000, 1:10000) and 25μl of each dilution spread on a quartile of a Columbia blood agar plate (CBA, Oxoid) before being incubated for 24 hours aerobically at 37°C to enumerate live bacteria. The organoids were washed a further 3 times in PBS before being lysed with 1% Triton-X100 in PBS for 10 minutes at RT. The lysate was added to CBA plates and incubated as above to detect intracellular bacteria. All experiments were completed in triplicate.

### Analysis of shed epithelial cells

Patient samples were collected with informed consent and analyzed in accordance with a protocol approved by the East Central London Regional Ethics Committee (REC1) (Ref: 11/H0721/7).

Experimentally infected organoid supernatants, or patient urine specimens, were collected, cytocentrifuged on to glass slides and fixed and stained prior to imaging^54^. Briefly, 80μl of supernatant or patient urine was cytocentrifuged using a Shandon Cytospin 2 cytocentrifuge at 800rpm (≈75g rcf) for 5 minutes. The cellular deposit was circumscribed with a ImmEdge pen (Vector Laboratories) before fixing, staining and mounting as above.

### Imaging and Analysis

We performed epi-fluorescence microscopy on an Olympus CX-41 upright microscope, and confocal laser scanning microscopy on Leica SP5 and SP2 microscopes. Images were processed and analysed using Infinity Capture and Analyze V6.2.0, ImageJ 1.50h^73^ and the Leica Application Suite, Advanced Fluorescence 3.1.0 build 8587 Software.

TEM was conducted using a Jeol 1200-Ex digital image capture system with a side mount 2Kv AMT camera. SEM was performed using a Jeol JSM-5300 fitted with a Semafore digital image capture system.

The proportion of the HBLAK organoids exhibiting evidence of differentiation was calculated by analyzing low-power SEM micrographs (N=3) using ImageJ automatic thresholding and measure tools^73^. The number of bacteria per cell was calculated using nearest neighbour 3D connectivity analysis with the ImageJ Object counter3D plugin^73,74^. The DAPI channel of 3D laser scanning confocal constructs were analysed using Differential Voxel Filters allowing the enumeration of mammalian nuclei and bacteria.

### Statistical analysis

Data were analysed using IBM SPSS Statistics version 24. Non-parametric Friedman tests of differences among repeated measures were performed due to non-normal distributions. 3 experimental replicates were performed for statistical testing.

## Data availability

All data generated or analysed during this study are included in this published article.

## Acknowledgements

We thank the MS Society for their generous support of this work (grant ref. 986), Peter Girling for helpful discussion, and Anthony Kupelian, Jakob Møller-Jensen and Paola Bonfanti for critical comments on the manuscript. We are also grateful to the Royal Free Elecron Microscopy Unit and to Linda Collins for expert technical assistance.

## Author contributions statement

HH and JR conceived the experiments, HH and DD conducted the experiments, HH, JML and JR analysed the results, and HH and JR wrote the manuscript. All authors reviewed the manuscript.

